# Nitric Oxide Synthase inhibition counteracts the stress-induced DNA methyltransferase 3b expression in the hippocampus of rats

**DOI:** 10.1101/2020.08.06.240374

**Authors:** Izaque de Sousa Maciel, Amanda Juliana Sales, Plinio C Casarotto, Eero Castrén, Caroline Biojone, Sâmia R. L. Joca

## Abstract

It has been postulated that activation of NMDA receptors (NMDAr) and nitric oxide (NO) production in the hippocampus is involved in the behavioral consequences of stress. Stress triggers NMDAr-induced calcium influx in limbic areas, such as the hippocampus, which in turn activates neuronal NO synthase (nNOS). Inhibition of nNOS or NMDAr activity can prevent stress-induced effects in animal models, but the molecular mechanisms behind this effect are still unclear. In this study, cultured hippocampal neurons treated with NMDA or dexamethasone showed increased of DNA methyltransferase 3b (DNMT3b) mRNA expression, which was blocked by pre-treatment with nNOS inhibitor n^ω^-propyl-L-arginine (NPA). In rats submitted to the Learned Helplessness paradigm (LH), we observed that inescapable stress increased of DNMT3b mRNA expression at 1h and 24h in the hippocampus. The NOS inhibitors 7-NI and aminoguanidine (AMG) decreased the number of escape failures in LH, and counteracted the changes in hippocampal DNMT3b mRNA induced in this behavioral paradigm. Altogether, our data suggest that NO produced in response to NMDAr activation following stress upregulates DNMT3b in the hippocampus.

## 1. Introduction

Exposure to unpredictable and uncontrollable stress triggers maladaptive emotional and behavioral responses associated with the development of neuropsychiatric disorders, such as post-traumatic stress disorder, anxiety and major depressive disorder (MDD) (Ulrich-Lai & Herman, 2009; Musazzi *et al.*, 2013; Sousa, 2016; Nava *et al.*, 2017). Dynamic regulation of gene expression is a critical component of adaptation to stress and engages epigenetic mechanisms, a process to alter gene expression without changing the sequence of DNA nucleotides (Mahgoub & Monteggia, 2013; Herre & Korb, 2019). DNA methylation is one of the main epigenetic mechanisms which is central to the neurobiology of MDD (Mahgoub & Monteggia, 2013; Matrisciano et al., 2016). The process involves the enzymatic addition of a methyl radical to cytosine-phosphate-guanine (CpG) group through DNA methyltransferases (DNMTs: 1, 2, 3a, 3b and L), favoring chromatin condensation and gene repression (Okano et al., 1999; Weber & Schübeler, 2007; Elliott et al., 2010; Feng et al., 2010; Mahgoub & Monteggia, 2013). The DNMT1 maintains DNA methylation during replication and the isoforms DNMT3a or DNMT3b are associated with *de novo* DNA methylation, inducing new patterns of methylation on CpG sequence (Bird, 1986; Okano et al., 1999; Elliott et al., 2010; Feng et al., 2010; Mahgoub & Monteggia, 2013).

Exposure to stress is associated with changes in DNMTs expression and DNA methylation in the promoter region of many genes relevant to neuronal function and neuroplasticity in cortical and limbic brain regions (Vialou et al., 2013; Nagy et al., 2017). Changes in DNA methylation and DNMT levels have also been described in blood cells of depressed patients, which are sensitive to chronic antidepressant treatment (Higuchi et al., 2011; Gassen et al., 2015). Similarly, chronic antidepressant treatment attenuates DNA methylation and DNMTs levels both *in vivo* and *in vitro* (Zimmermann et al., 2012; Vialou et al., 2013; Sales & Joca, 2018).

Direct inhibition of DNA methylation promotes antidepressant-like effects associated with increased levels of brain-derived neurotrophic factor (BDNF) in the hippocampus (Sales *et al.*, 2011; Sales & Joca, 2016). A growing body of evidence reports increased DNA methylation of the *Bdnf* gene promoter region in the cortex and hippocampus of rodents submitted to stress, resulting in decreased expression (Schmidt & Duman, 2007; Adachi et al., 2008; Roth & Sweatt, 2011; Roth et al., 2011). BDNF plays a key role in several physiological processes in the developing and mature brain, such as neurogenesis, neuroprotection and regeneration, as well as reinforcing short- and long-lasting synaptic interactions (Kowiański et al., 2018). Decreased levels of BDNF is associated with impaired synaptic transmission often described in stressed animals and MDD patients (Duman & Duman, 2015; Castrén & Antila, 2017; Kowiański *et al.*, 2018).

Stress exposure leads to HPA axis activation and release of glucocorticoids, which facilitates glutamate release in limbic brain regions, a mechanism that is often associated with functional and morphological impairments induced by exposure to uncontrollable stress (Sousa & Almeida, 2012; Duman & Duman, 2015; Belleau et al., 2019). Activation of both glucocorticoid and NMDA receptors can regulate the activity of the enzymes involved in epigenetic mechanisms, thus regulating gene expression in response to stress (Chandramohan et al., 2008). Activation of NMDAr triggers the synthesis of nitric oxide (NO) in the brain, which can rapidly diffuse across membranes and modulate several inter- and intracellular processes (Prast & Philippu, 2001; Guix et al., 2005). There are three major NO synthases isoforms: nNOS (neuronal; NOS1) and eNOS (endothelial; NOS3), which are constitutively expressed and Ca2+-calmodulin-dependent; and iNOS (inducible; NOS2), which is mostly expressed under inflammatory or immunological stimuli (Guix et al., 2005; Amitai, 2010). Excessive NO production induced by stress is neurotoxic and predisposes to behavioral changes and MDD (Bishop & Anderson, 2005; Calabrese et al., 2007). Accordingly, administration of nNOS or iNOS inhibitors attenuates stress-induced NO increase in the CNS, and induces antidepressant-like effects (Joca & Guimarães, 2006; Montezuma *et al.*, 2012; Hiroaki-Sato *et al.*, 2014; Stanquini *et al.*, 2017). Accumulating evidence indicates that NO-induced effects might be related to its ability to regulate BDNF-TrkB signaling and neuronal plasticity in the mature brain (Biojone et al., 2015).

NO can also regulate activity and expression of chromatin-modifying enzymes, thus posing an important role for NO in modulating epigenetic mechanisms (Campos *et al.*, 2007; Katayama *et al.*, 2009; Vasudevan *et al.*, 2016; Socco *et al.*, 2017). However, it is not known if NO release during exposure to stress increases the DNMTs expression and DNA methylation in the brain.

Thus, we hypothesized that the antidepressant-like effect of NOS inhibitors is associated with a decrease in DNA methylation. To this aim, we assessed the effect of NOS inhibitors in the expression of DNA methyltransferases in cultured hippocampal cells exposed to stress mediators and rats exposed to inescapable foot shocks.

## 2. Methods

### 2.1. Animals

Male Wistar rats (8 weeks old, weighing 270-300g) were obtained from the animal facility of the University of São Paulo/USP Ribeirão Preto. The animals were kept undisturbed for one week before the beginning of the experiments, except for regular cage cleaning, in groups of four per cage (41 × 34 × 16cm) at 24±1°C under standard laboratory conditions (12h light/12h dark, lights on at 06:30 a.m.) with free access to food and water. During the experimental procedure, the animals were kept isolated (30 × 20 × 13cm) with free access to food and water (Sales & Joca, 2018). The total number of animals used in this study were 170. All procedures were conducted in accordance with the Brazilian Council for Animal Experimentation (COBEA), which comply with international laws and policies. The protocols were approved by the local Ethical Committee (protocol number 15.1.285.60.5.), and all efforts were made to minimize animal suffering and to reduce the number of animals used. The experiments were conducted between 08:00-17:00 with randomization of the experimental groups throughout the day.

### 2.2. Pharmacological Treatment

The animals received intraperitoneal injections of aminoguanidine - AMG (30mg/kg, Sigma-Aldrich (#396494), St. Louis, MO, USA), 7-nitroindazole - 7NI (60mg/kg, Sigma-Aldrich (#N7778), St. Louis, MO, USA) or vehicle (NaCl 0.9% for AMG; polyethylene glycol and NaCl 0.9% - 1:1 - for 7-NI). The drugs were injected immediately after the LH pretest session and repeated once a day for 7 days. The last injection was performed 1h before the behavioral test or euthanasia. The dose and schedule of treatment were based in literature (Joca *et al.*, 2003; Stanquini *et al.*, 2017; Sales & Joca, 2018). For *in vitro* experiments, the nNOS inhibitor n^ω^-propyl-L-arginine (NPA, 100nM, Sigma–Aldrich, #SML2341) was administered 30 minutes before the stimulus with NMDA (30μM for or 1h; Sigma–Aldrich, #M3262), dexamethasone (1 μM for 1h or 24h, Sigma–Aldrich, #D4902) or L-arginine (0.5mM for 1h, Sigma–Aldrich, #A5006). All concentrations and schedules of treatment were based on previous studies in the literature (Sandoval *et al.*, 2011; Anacker *et al.*, 2013; Xiong *et al.*, 2014).

### 2.3. Learned helplessness test

The learned helplessness (LH) model is based on the exposure to inescapable footshocks, which leads to increased number of escape failures in a later learning associative task (Maier & Watkins, 2005; Maier & Seligman, 2016). Increased circulating glucocorticoid levels and glutamate release, as well as increased synthesis and release of nitric oxide (NO) have also been described in this paradigm (Prast & Philippu, 2001; Joca *et al.*, 2003; Popoli *et al.*, 2011; Nava *et al.*, 2017; Stanquini *et al.*, 2017; Sales & Joca, 2018). Repeated treatment with antidepressants (Joca *et al.*, 2003) and nNOS inhibitors (Stanquini *et al.*, 2017) attenuates the number of escape failures during the LH test, considered a reliable antidepressant-like effect.

The behavioral experiment consisted of two sessions (pretest and test), as previously described (Stanquini *et al.*, 2017). During the pretest, the animals were randomly assigned to one of the following conditions: stressed group (IS – inescapable shocks) or habituated group (hab – no-shocks). The animals in the IS group were individually placed into the automated shuttle boxes (307 × 333 × 540mm, Model EP 11, Insight Scientific Equipment), with two compartments of equal size separated by a wall with a central open door (communicating the two compartments), and forty inescapable foot shocks (0.4mA, 10s duration) were given in a variable schedule with a mean interval of 60s (range from 30–90s) through the metal grid floor. The habituated group was exposed to the same apparatus for 30 min, but no shock was delivered. After six days, both groups were submitted to the test session (T). The test consisted of 30 escapable foot shocks (0.4mA, 10s duration, 30–90s interval), which were preceded by a tone (60dB, 670Hz) that started 5s before each shock and lasted until its end. Animals could prevent or interrupt the shock delivery by either crossing to the opposite side of the box during sound presentation (avoidance) or by crossing during its presentation (escape). The absence of any of these behaviors was considered escape failure. The number of crossings during the intervals between shocks, in the test session, was used as an index of locomotor activity (Geoffroy & Christensen, 1993; Stanquini *et al.*, 2017; Sales & Joca, 2018).

### 2.4. Cell culture

Primary cultures of hippocampal cells from E17-E18 rat embryos were prepared as detailed in literature (Sahu *et al.*, 2019). Briefly, hippocampus of E18 rat embryos were dissected and suspended cells were seeded on poly-L-lysine coated 24-well plates (250,000 cells/well, 1.9cm^2^) in Neurobasal medium supplemented with B27, 1% penicillin/streptomycin, 1% L-glutamine. The medium was changed weekly, and the cells were treated and lysated between DIV 8-10.

### 2.5. Gene expression analysis by quantitative real-time RT-PCR (RT-qPCR)

Total RNA was extracted using Trizol® reagent (from Life technologies) according to manufacturer’s instruction and quantified by spectrophotometer (Nanodrop 2000C spectrophotometer, Thermo Scientific). The cDNA was synthesized from 2 μg of total RNA using Maxima first strand cDNA synthesis kit for RT-qPCR with dsDNase (Thermo Scientific) according to the manufacturer’s instruction. Two controls of reaction were performed: reverse transcriptase minus (RT-) negative control, which contains all reagents for the reverse transcription reaction except the Maxima Enzyme Mix; and the non-template control (NTC), which contains all reagents for the reverse transcription reaction except the RNA template. The primers used to amplify specific cDNA transcripts are listed in table 1 (Karpova *et al.*, 2010). Quantitative PCR was performed using Thermo Scientific Maxima SYBR Green qPCR Master Mix (2x). The reaction was done in a volume of 20μl (10μl SYBR Green Master Mix 2x; 4μl water – nuclease free; 0,5μl of primers (forward and reverser, 0.25 μM, final concentration) and 5μl of cDNA. The DNA amplification reaction was assayed in triplicates in the Bio-Rad C1000TM Thermal Cycle. The qPCR condition was set at 95°C for 10 min (denaturation) and 44 cycles at 95°C 15s (denaturation); temperature of annealing (mT), depending on the primers used for 30s and 72°C for 30s (extension). At the end of the cycling protocol, a melting-curve analysis was included, which showed a single peak in all experiments. *Hprt1* gene expression was not modified by any of the treatments, and it was then chosen as housekeeping gene for later relative quantitations. The relative expression levels for each target gene was calculated by the 2^−(ΔΔCT)^ method (Livak & Schmittgen, 2001). The results were expressed as % from vehicle-treated groups.

**Table 1.**
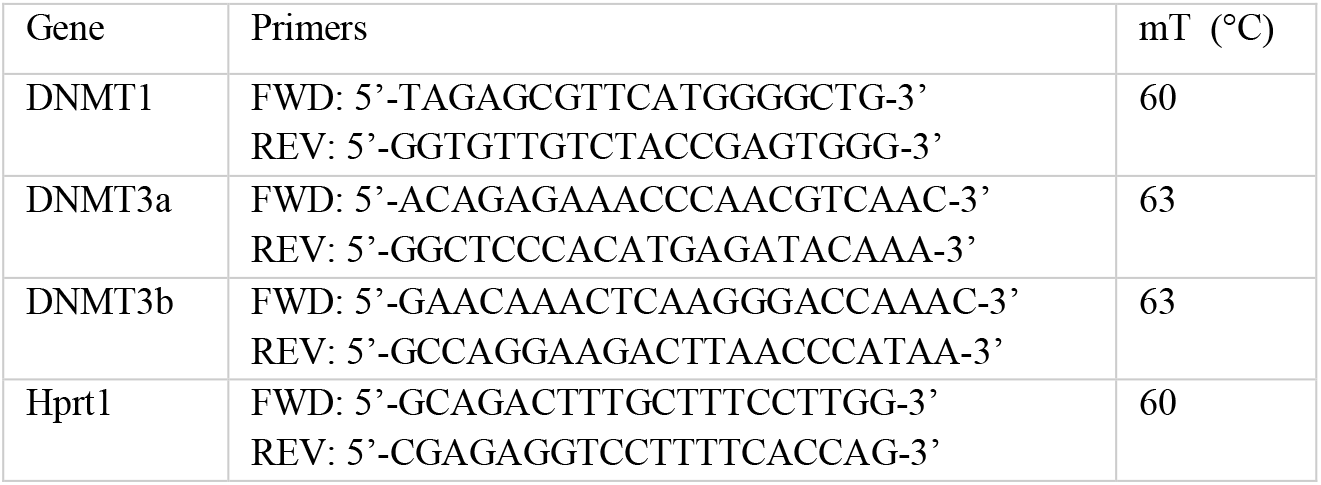
Sequence of primers.

### 2.6. Experimental Design

#### 2.6.1. NMDA-nNOS/NO signalling effects in DNA methyltransferase mRNA

Hippocampal cell cultures (DIV 8-10) were pre-treated with NPA (100nM) followed, 30min later, by NMDA (1μM). One hour after, mRNA was extracted and the DNMT1, DNMT3a and DNMT3b mRNA were determined by RT-qPCR as described. Another batch of cells was pre-treated with NPA (100nM) followed by L-arginine (0.5 mM). The mRNA was extracted, one hour after the last drugs administration and the DNMT3b mRNA was assessed by RT-pPCR.

#### 2.6.2. Effect of dexamethasone in DNMT3b mRNA expression

In another experimental set, hippocampal cell cultures (DIV 8-10) were pre-treated with NPA (100nM) followed by dexamethasone (1μM). One or 24 hours after the last drug administration the mRNA was extracted as described and the DNMT3b mRNA were determined by RT-qPCR.

#### 2.6.3. Antidepressant-like effect of NOS inhibitors

The behavioral effects of NOS inhibitors were assessed using the learned helplessness paradigm as described. The drug effect was evaluated in the stressed group (exposed to inescapable foot shocks) and the habituated group (exposed to shuttle box without foot shocks). The experimental number for each group was based as calculated in our previous publication (Stanquini *et al.*, 2017). Each experimental session included at least 2 animals of each treatment group and the experiment was repeated 5-6 times, always including animals of the different treatment conditions. Immediately after PT session the rats were injected with AMG (30 mg/kg, ip) or vehicle (saline 0.9%, ip) and once a day for the following 6 days. The last injection was performed 1h before the behavioral test. Another group of rats was treated with 7NI (60 mg/kg, ip) or vehicle (polyethylene glycol and NaCl 0.9% - 1:1, ip) under the same protocol described. On the seventh day, the number of failures, latency to escape and inter-trail crossing were automated assessed.

#### 2.6.4. Increase of DNMT3b mRNA in the hippocampus of rats exposed to inescapable foot shocks

Based on the *in vitro* results, an independent group of animals were exposed to inescapable footshock (stressed group) and euthanized 1h, 24h or 7 days later for mRNA extraction from the hippocampus (vHPC and dHPC) as described, and the DNMT3b mRNA was determined. In another set of experiments, the animals were treated with AMG or 7NI immediately after PT session, followed by euthanizing 1h after the drug administration. Another other experimental group was exposed to two injections, the first immediately and the second 23h after the PT, followed by euthanasia 1h after the last drug administration. The RNA extraction and RT-qPCR were evaluated as described.

### 2.7. Statistical analysis

Discrete variables, such as the number of failures, avoidance, and intertrial crossings in LH experiments were analysed by non-parametric tests (Kruskal-Wallis), followed by Dunn’s multiple comparison whenever appropriate. The *in vitro* experiments were analyzed by two-way ANOVA followed by Bonferroni’s test, with stress and pharmacological treatment as main factors, or Student’s t-test. Results of RT-PCR from *in vivo* experiments were analyzed by one-way ANOVA followed by Newman-Keuls’ test. P values below 0.05 were considered statistically significant. All data used in the present manuscript is available under CC-BY license in FigShare (DOI:10.6084/m9.figshare.12424349).

## 3. Results

### 3.1. NOS inhibitor NPA attenuated DNMT3b increase in hippocampal primary cell culture challenged with NMDA or dexamethasone

To investigate the molecular mechanisms associated with NO effects on DNA methylation *in vitro*, we used hippocampal cell cultures. In order to simulate stress effects on the cells, we used the glucocorticoid receptor agonist dexamethasone, or NMDA to activate NMDAr. Previous studies indicate stress triggers increased glutamate release in the brain and the activation of NMDAr and nNOS/NO signaling in hippocampus (Popoli *et al.*, 2011; Sandoval *et al.*, 2011; Musazzi *et al.*, 2013; Joca *et al.*, 2015; Wegener & Joca, 2016). Thus, we exposed cultured hippocampal cells to NOS inhibitor NPA (100nM/30min), followed by NMDA (30μM/1h), figure 1A. The pretreatment with NPA 100nM prevented NMDA-induced DNMT3b mRNA increase (interaction, F_1,32_= 4.851, p= 0.03, N= 9; figure 1B). However, the DNMT1 (interaction, F_1,20_= 0.02, p= 0.90, N= 6; figure 1D) or DNMT3a (interaction, F_1,20_= 0.08, p= 0.78, N= 6; figure 1E) mRNA amount were not affected by the NMDA or NPA treatments. To evaluate if nNOS/NO activation could directly induce an increase of DNMT3b mRNA in hippocampal primary cell culture, we assayed the DNMT3b mRNA after the treatment with the NOS substrate L-arginine (0.5mM/1h) (Xiong *et al.*, 2014). L-arginine increased the DNMT3b mRNA, which was attenuated by NPA pretreatment (interaction, F_1,20_= 13.35, p= 0.0009, N= 6; figure 1C).

**Figure 1:**
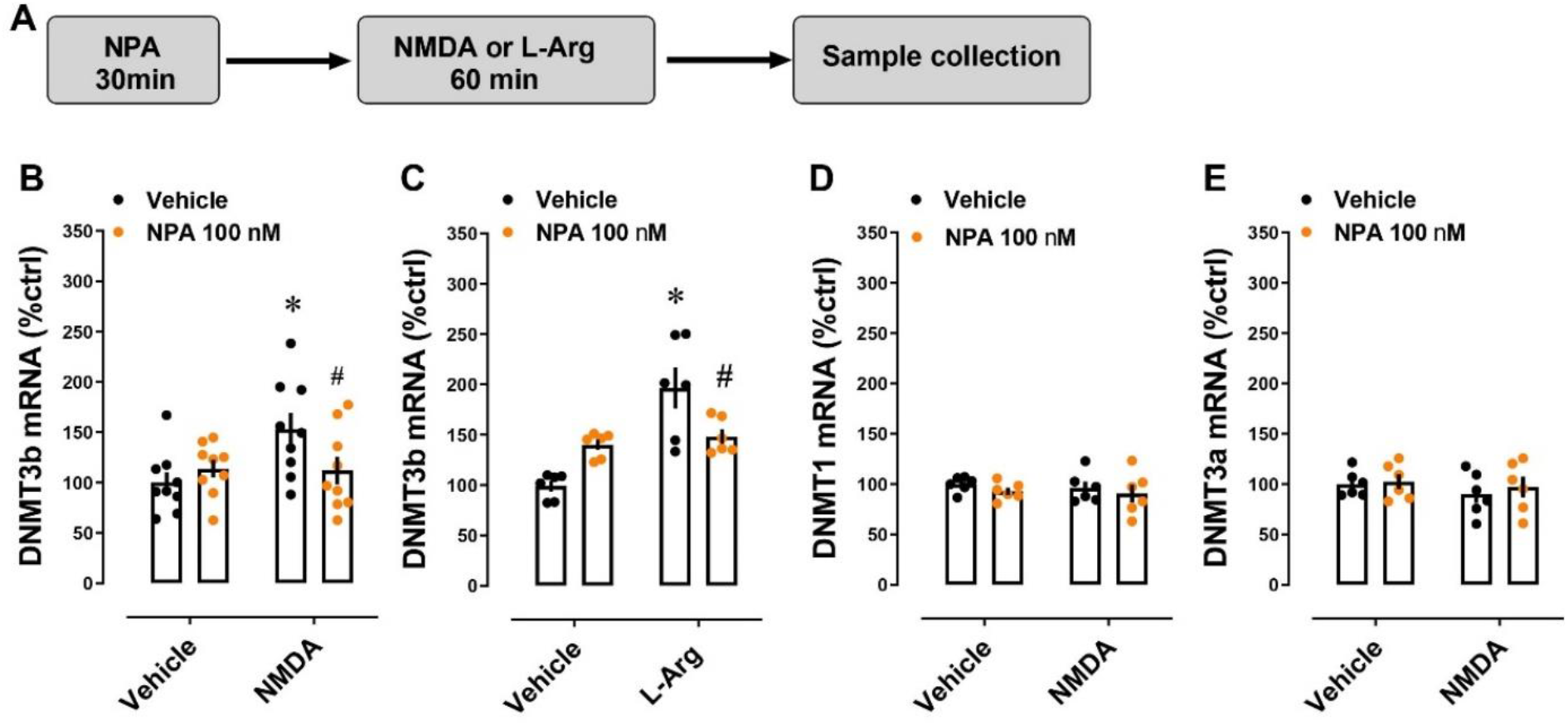
nNOS inhibitor NPA prevented the increase in DNMT3b mRNA in neuronal hippocampal cell culture challenged with NMDA or L-arginine: (A) Timeline of experimental approach. The previous treatment with NPA (100nM/30min) attenuates the increase of DNMT3b mRNA induced by (B) NMDA (30μM/1h) or (C) L-arginine (0.5mM/1h). No interaction was observed for the amount of (D) DNMT1, (E) DNMT3a after challenge with NMDA, N=6-9. Each column represents the mean ± SEM. *p<0.05 from vehicle/vehicle group; #p<0.05 from the NMDA/vehicle or L-Arg/vehicle groups.

Based on the role of corticosterone/dexamethasone on the activation of NMDA receptors in hippocampus (Takahashi *et al.*, 2002), we also tested whether dexamethasone could induce similar changes on DNMTs mRNA amount *in vitro*. As expected, dexamethasone (1μM/24h) increased DNMT3b mRNA, which was attenuated by pretreatment with NPA (100nM/30min) (interaction: F_1,20_= 20.29; p= 0.0002, N= 6; figure 2C). However, the DNMT3b mRNA was not changed after only 1h incubation with dexamethasone (F_1,8_= 0.1225; p= 0.74, N= 3; figure 2B) or by pretreatment with NPA (interaction: F_1,8_= 1.066; p= 0.33, N = 3; figure 2B). Moreover, the DNMT1 (1h: t= 1.644, df= 4, p = 0.175; figure 2D and 24h: t = 1.122, df= 10, p= 0.288; figure 2F) or DNMT3a (1h: t= 0.55, df= 4, p= 0.61; figure 2E and 24h: t= 0.35, df= 10, p= 0,73; figure 2G) mRNA did not change after dexamethasone treatment.

**Figure 2:**
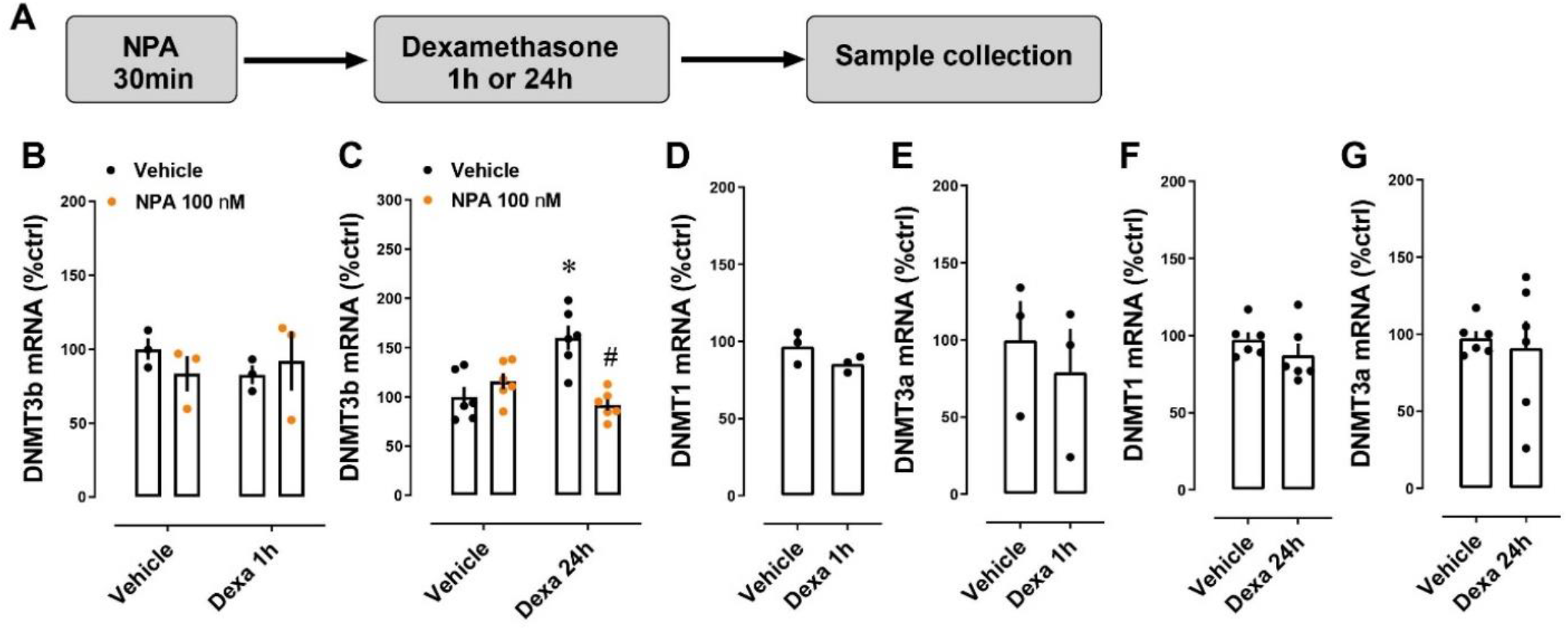
nNOS inhibitor modulated the DNMT3b mRNA expression in hippocampal cell culture challenged with dexamethasone: (A) Timeline of experimental approach. Relative amount of DNMT3b (B and C), DNMT1 (D and F), and DNMT3a (E and G) mRNA after pretreatment with NPA (100nM/30min) followed by dexamethasone (1μM) for 1h or 24h, N=3-6. Each column represents the mean ± SEM. *p< 0.05 from the vehicle/vehicle group; #p<0.05 from the dexamethasone/vehicle group.

### 3.2. NOS inhibitors attenuated the depressive-like behavior and the increase of DNMT3b mRNA in rats submitted to the LH

Repeated treatment with AMG (Kruskal-Wallis, H= 25.64, df= 3; p= 0.03; N= 11-13) or 7NI (Kruskal-Wallis, H(3)= 15.65, df= 3; p= 0.02; N= 10-13), figure 3B-C, attenuated the increase in escape failures, as well the latency time to escape/avoid shocks (AMG, interaction: F_1,45_ = 4.924; p = 0.03; N = 11-13, figure 3D or 7NI, interaction: F_1,40_= 6.45; p= 0.01; N= 10-13, figure 3E) in rats submitted to LH paradigm. Neither stress nor pharmacological treatment changed the locomotor activity assessed by intertrial crossing (H= 1.664, df = 3; p= 0.99; N= 11-13 and H= 2.731, df = 3; p= 0.99; N= 10-13, for AMG and 7-NI, respectively; figure 3F and 3G).

**Figure 3:**
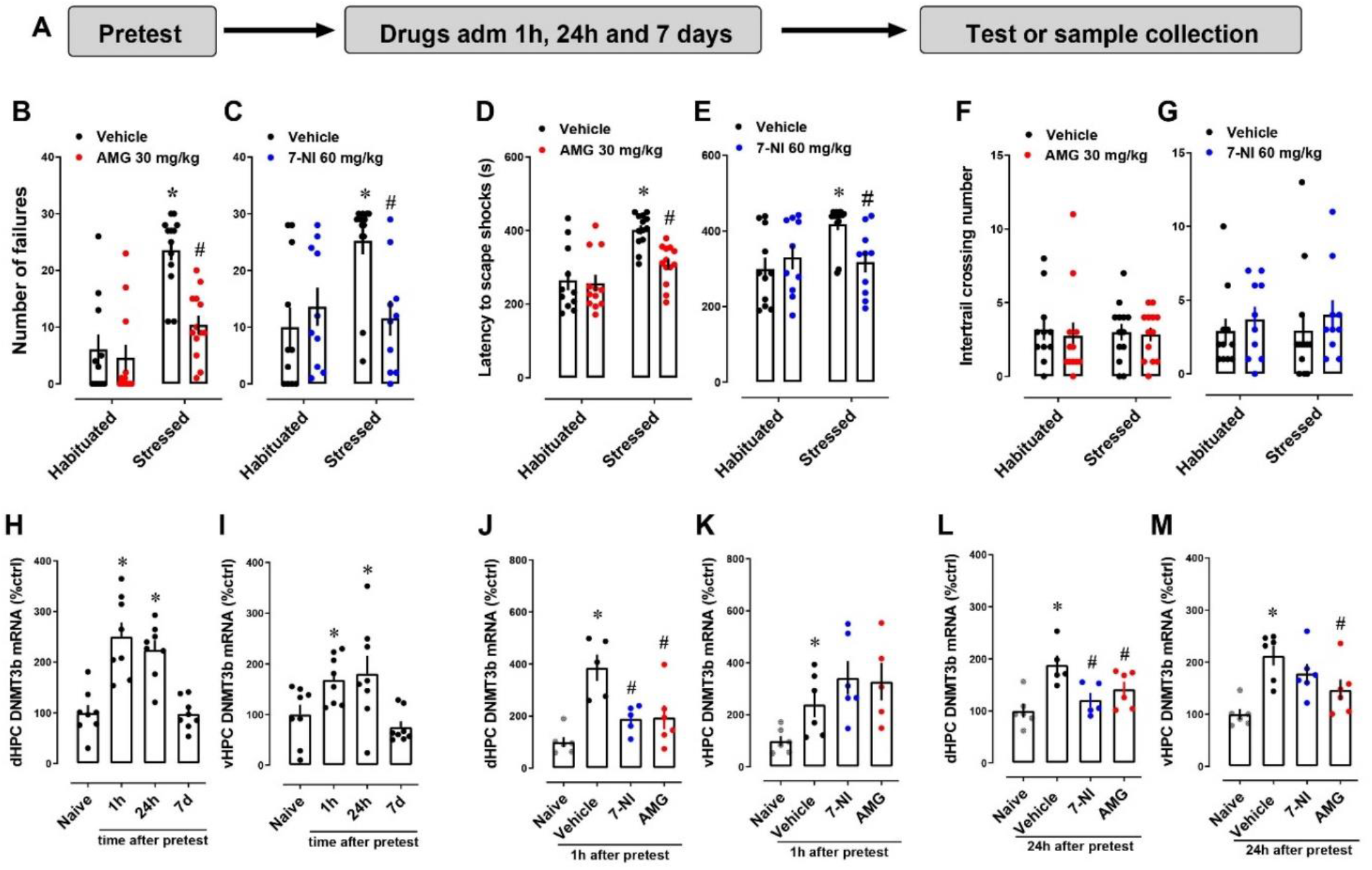
Antidepressant effect of NOS inhibitors is associated with a decrease in DNMT3b mRNA in the hippocampus. (A) Timeline of experimental approach. The animals were submitted to inescapable foot shocks (IS, stressed) or habituated (hab, non-stressed) in a shuttle-box. Immediately after IS, the rats were treated with AMG (30mg/kg); 7-NI (60mg/kg) or vehicle, once a day for up to 7 days (N= 13-14). One hour after the last injection, the rats were submitted to the LH test session or euthanized for hippocampus dissection. The administration of AMG or 7-NI attenuated the effect of IS in (B and C) number of failures, (D and E) latency time to escape/avoid shocks but no alterations were observed in (F and G) locomotor activity assessed by intertrail crossing. Independent groups of animals were submitted to IS and the brains were dissected after 1h, 24h or 7 days (H and I), N=6-8. To assess the effect of NOS inhibitor in DNMT3b mRNA amount, immediately after pretest session (IS) the rats were treated with one i.p injection (1h, J and K) or 2 i.p, injections, one immediately after pretest session and another 1h before the brain dissection (24h, L and M). Data are expressed as mean ± S.E.M. * p< 0.05 from hab/vehicle or naive groups; #p< 0.05 from stressed/vehicle group. PT: pretest; IS: inescapable footshock; Hab: habituated, non-stressed; AMG: aminoguanidine; 7-NI: 7-nitroindazole.

In order to assess the effect of inescapable and uncontrollable foot shock on the DNMT3b mRNA in the hippocampus, we assayed the mRNA levels at 1h, 24h and 7 days after the pretest session. Exposure to inescapable and uncontrollable footshocks in pretest session induced an increase of DNMT3b mRNA in dHPC (F_3,28_= 17.16; p= 0.0001, N= 8, figure 3H) and in vHPC (F_3,28_ = 5.269; p = 0.005, N= 8, figure 3I) after 1h and 24h. However, in both hippocampal regions, the amount of DNMT3b mRNA returned to basal 7 days after exposure to the shocks (figure 3H,I). In dHPC, the treatment with both NOS inhibitors (AMG or 7NI) prevented the increase of DNMT3b mRNA at 1h (F_3,18_= 10.06; p= 0.0004, N = 5-6; figure 3J) and 24h (F_3,18_= 6.302; p= 0.004, N= 5-6; figure 3L). On the other hand, in vHPC, the treatment with AMG attenuated the stress-induced increase of DNMT3b mRNA at 24h (F_3,20_= 7.635; p = 0.001, N= 6; figure 3L), but did not change the mRNA quantity at 1h (F_3,19_= 4.386; p= 0.02, N= 5-6; figure 3K). In both time points assayed (1h and 24h), the treatment with 7NI did not inhibit the increase of DNMT3b mRNA (1h: F_3,19_= 4.386; p= 0.02, N= 6 and 24h: F_3,20_= 7.635; p= 0.001, N= 6; figure 3J,K).

## 4. Discussion

The present study describes the effect of NOS inhibitors on DNMT3b mRNA expression induced by stress. Challenging cultured hippocampal neurons with mediators of the stress response (NMDAr agonist or the glucocorticoid dexamethasone) increased DNMT3b mRNA expression. Rats exposed to inescapable and uncontrollable foot shocks displayed learned helplessness behavior, characterized by an increase in the number of failures to escape foot shocks in the test session. Such phenotype was associated with increased DNMT3b mRNA expression in the hippocampus, and in both *in vitro* and *in vivo* approaches, the pre-treatment with NOS inhibitors attenuated the stress-induced increase in DNMT3b mRNA expression.

Acute stress triggers a complex response involving the synthesis and release of mediators such as corticoids, glutamate and NO, which can acutely alter gene expression as well as induce long-term molecular and behavioral changes (Popoli et al., 2011; Joca et al., 2019). The increase in NO and/or reactive oxygen species (ROS) can directly impact gene expression (Riccio *et al.*, 2006; Lee *et al.*, 2009; Nott & Riccio, 2009). For instance, oxidative stress can increase the activity of specificity protein (Sp), a transcription factor involved in DNMT expression (Ryu *et al.*, 2003; Jinawath *et al.*, 2005; Lin & Wang, 2014; Chuang *et al.*, 2017), and NO facilitates cGMP, PKG, and ERK signaling cascades, involved in the activation of CREB, which in turn modulates the expression of several genes (Lu *et al.*, 1999; Gallo & Iadecola, 2011).

Little attention has been given to the role of NO on DNA methylation. Some sparse evidence indicates that cellular stress increases NO production, followed by DNMT3b and DNMT1 expression (Campos *et al.,* 2007; Huang *et al.*, 2012). Also, the NO donor (Sodium Nitroprusside-SNP) increases the DNMT activity and decreases fragile X retardation 1 (FMR1) gene expression in Jurkat T cell (Hmadcha et al., 1999). We observed that NPA prevented the increase in DNMT3b levels by NMDA or L-arginine in cultured hippocampal cells. Corroborating these findings, the non-selective NOS inhibitor (L-NAME) attenuates the DNMT expression, and the levels of 5-Methylcytosine, indicative of global DNA methylation, in melanoma cells under stress (Campos *et al.*, 2007). However, in our conditions, NMDA significantly increased DNMT3b mRNA, but not DNMT3a or DNMT1 mRNA.

Increased activation of HPA-axis and corticoid receptors in the brain is involved in genomic and non-genomic changes induced by stress (McEwen *et al.*, 2016; van Bodegom *et al.*, 2017). As described above, 24h dexamethasone treatment in hippocampal cell culture increased DNMT3b mRNA levels, which was prevented by NPA. In line with our data, dexamethasone treatment increases DNA methylation in the promoter region of the *crh* gene. Such mechanism is associated with the formation of a repressor complex consisting of glucocorticoid receptor (GR) and DNMT3b in rat hypothalamic IVB cells (Sharma *et al.*, 2013). Also, cultured primary cortical neurons treated with corticosterone (10 μM) show increased levels of DNMT1, DNMT3a and DNMT3b mRNA, which are abolished by co-treatment with GR antagonist mifepristone (20 μM) (Urb *et al.,* 2019). Thus, it is possible to assume that the genomic effects induced by GR activation may lead to increased DNMTs expression and DNA methylation in neurons. In addition, membrane glucocorticoid receptors facilitate and sustain calcium influx through NMDAr, resulting in Ca^2+^ neurotoxicity in hippocampal neurons (Takahashi et al., 2002). Together, these data suggest that DNMT mRNA expression is stimulated by stress mediators, and NO is a key mediator of DNMT3b mRNA expression triggered by glucocorticoids and NMDA-NOS/NO signaling.

NO has been involved in many signaling pathways related to stress and MDD, as well as to the antidepressant activity of classical drugs (Wegener et al., 2003; Joca & Guimarães, 2006; Wegener & Volke, 2010; Hiroaki-Sato et al., 2014). Corroborating the results from the present study, pharmacological inhibition of NOS induces antidepressant-like effects in the learned helplessness paradigm (Stanquini et al., 2017) and other models, such as forced swimming test (Ferreira et al., 2012; Montezuma et al., 2012; Sales et al., 2017) and unpredictable chronic stress (Yazir et al., 2012). Furthermore, activation and expression levels of NOS is attenuated by treatment with antidepressant drugs (Finkel et al., 1996; Wegener et al., 2003; Dhir & Kulkarni, 2007, 2011; Zomkowski et al., 2010; Zhou et al., 2011; Bollinger et al., 2017). Accordingly, MDD patients treated with selective serotonin reuptake inhibitors (SSRI) present a decrease of nitrate levels in the blood (Finkel et al., 1996), and rodents treated with SSRIs exhibit a reduction in the NOS activity in the brain (Krass et al., 2011).

Here, the repeated administration of NOS inhibitors counteracted the behavioral effects of uncontrollable stress, similarly to the treatment with classical antidepressant imipramine (Stanquini et al., 2017). Moreover, the behavioral effects of NOS inhibitors were associated with attenuation of the DNMT3b mRNA expression, which peaked in the hippocampus from 1h to 24h after stress. To our knowledge, this is the first evidence that the antidepressant-like effect of NOS inhibitors could be associated with the attenuation of stress-induced DNMT3b/DNA methylation.

Although acute treatment with NOS inhibitors is not sufficient to rescue the behavior change in LH (Stanquini *et al.*, 2017), some studies suggest that, at the molecular level, acute administration of NOS inhibitors in rodents can modulate transcription factors, as well as gene expression (Joca *et al.*, 2007; Nott & Riccio, 2009; Zhou *et al.*, 2011). Our data indicates that the pharmacological effect of AMG or 7NI on DNMT3b mRNA expression could be the result of acute inhibition of transcription factors activity mediated by NO in the hippocampus of rats exposed to inescapable foot shocks. Thus, the antidepressant effect induced by NOS inhibition, *i.e.* the facilitated adaptation to stress, could be the long-term consequence of the inhibition of DNMT3b/DNA methylation.

The hippocampus is highly susceptible to the effects of stress, seen as impaired function and changes in gene expression (Crosio *et al.*, 2003; Gray *et al.*, 2014). Previous data show that pharmacological or genetic inhibition of DNMT promotes antidepressant phenotype in rodents (Sales *et al.*, 2011; Morris *et al.*, 2016; Sales & Joca, 2016). Direct inhibition of DNMTs into the dorsal hippocampus induces similar effects in rats exposed to stress (Sales *et al.*, 2011), thus posing a central role for epigenetic mechanisms in this structure during stress. Also, stressed animals show an increase of global DNA methylation in HPC (Doherty et al., 2016; Sales & Joca, 2016), which is attenuated by antidepressant drugs treatment (Sales & Joca, 2016). Thus, it is possible to consider that, during stress, activation of corticoid receptors facilitates glutamate release and NMDA activation in the hippocampus, ultimately leading to NO synthesis. However, it is unclear how NO could regulate DNMT3b expression, but it is plausible to consider that it involves nuclear transcription factors, such as specificity protein Sp1, Sp3 or CREB, as proposed in figure 4. We speculate that the antidepressant-like effect of NOS inhibitors in rats exposed to LH test is associated with inhibition of mechanisms that control DNMT3b mRNA expression in hippocampus neurons.

**Figure 4:**
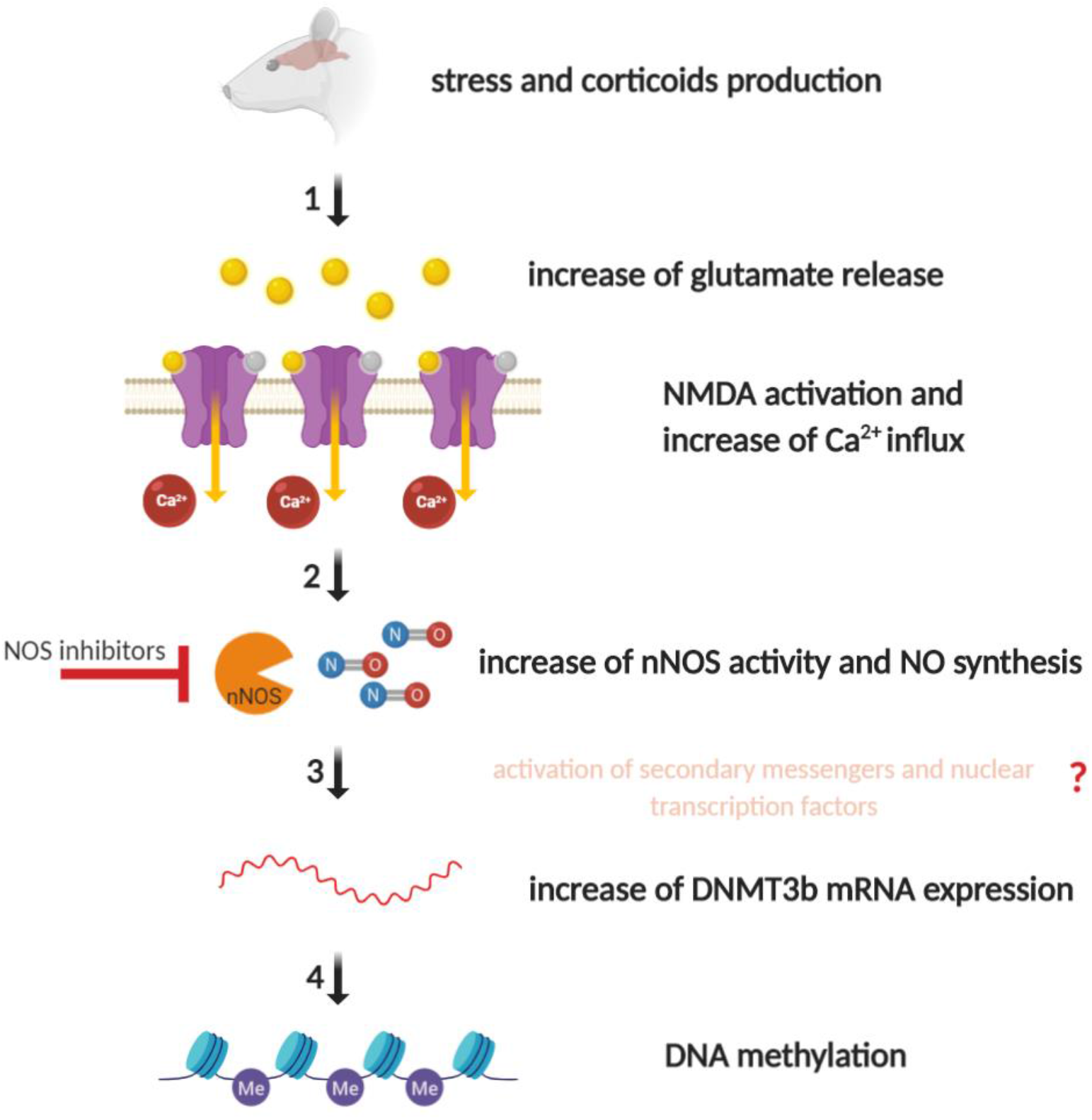
Summary of the hypothetical role of NO in DNA methylation induced by stress. The stress induced by uncontrollable and unpredictable foot shocks increases the release of glutamate in the hippocampus (1), which activates NMDA receptors and increases calcium influx. In hippocampus neurons, the increase of Ca^2+^ influx promotes nNOS activation and NO synthesis, which could increase the activity of secondary messengers and nuclear transcription factors, ultimately leading to increased DNMT3b expression (2). The NOS inhibitors mitigated the increase of DNMT3b during stress, which may prevent the DNA methylation in the hippocampus of rats (3 and 4).

Fanselow and Dong (Fanselow & Dong, 2010) published a comprehensive review about the behavioral, anatomical and patterns of gene expression studies showing a functional segmentation of hippocampus in three compartments (dorsal, intermediate, and ventral). At the molecular level, gene expression in the dHPC correlates with cortical regions involved in information processing (cognitive function), while in the vHPC this expression correlates with regions involved in the control of emotion and response to stress (amygdala and hypothalamus). According to these authors, the exact boundaries between these subregions are not clear and that the intermediate portion of HPC have distinct neuronal connectivity patterns, which influence gene expression (Fanselow & Dong, 2010). Furthermore, dHPC and vHPC are impacted differently by acute or chronic stress at genomic and proteomic level, as well as by pharmacological treatment (Diniz et al., 2016; Floriou-Servou et al., 2018). Data from our group showed that acute stress (FST) increases the phosphorylation of nNOS in dHPC, but not in vHPC of rats (Diniz et al., 2016). Interestingly, the local administration in dHPC of the nNOS inhibitor NPA immediately after the pretest session in FST, or 1h before the test session, decreases the immobility time in FST. On the other hand, in the vHPC, this compound decreases the immobility time only when infused 1h before the test (Diniz et al., 2016). These data are in line with the results from the present study, where both NOS inhibitors (AMG or 7NI) treatment decreased the DNMT3b mRNA expression on dHPC at 1h (one injection) and 24h (two injection) after stress. In vHPC, only two injections (1h and 23h after stress) of AMG, significantly decrease the DNMT3b mRNA expression. Together, these results indicate that dHPC and vHPC have different sensitivity to stress and to the pharmacological effect of NOS inhibitors.

In conclusion, we showed for the first time that NOS inhibitors may control DNMT3b mRNA expression in rat hippocampus. Our results suggest the participation of NO in the cellular pathways related to DNA methylation triggered by unpredictable and uncontrollable stress. Therefore, it is possible to assume that NOS enzyme is a putative target to control the consequences of hyperactivation of HPA-axis, DNMT/DNA methylation, and gene expression involved with the neurobiology of major depression.

## Abbreviations

NMDAr: NMDA receptor
NO: nitric oxide
nNOS: neuronal NO synthase
DNMTs: DNA methyltransferases
DNMT3b: DNA methyltransferase isoform 3b
NPA: n-ω-propyl-L-arginine
LH: Learned Helplessness paradigm
7NI: 7-Nitroindazole
AMG: aminoguanidine
MDD: major depressive disorder
CpG: cytosine-phosphate-guanine
BDNF: brain-derived neurotrophic factor
HPA axis: Hypothalamic-Pituitary-Adrenal axis
CNS: Central nervous System
TrkB: Tropomyosin receptor kinase B
L-arg: L-Arginine
IS: Inescapable Shocks
hab: habituated group
PT: pretest
T: test
dHPC: dorsal hippocampus
vHPC: ventral hippocampus
ROS: reactive oxygen species
Sp: specificity protein
cGMP: cyclic guanosine monophosphate
PKG: protein kinase G
ERK: extracellular signal-regulated kinase
CREB: cAMP responsive-element binding protein
SNP: Sodium Nitroprusside
FMR1: fragile X retardation 1
GR: glucocorticoid receptor

## Conflict of interest and funding

EC has received lecture fees from Janssen-Cilag. All other authors declare no conflict of interest. This study was supported by Brazilian research agencies FAPESP (#2015/25067-6, #2015/06271-1 and #2017/24304-0; São Paulo Research Foundation Brazil), CNPq (#306648/2014-8 and #304780/2018-9; National Council for Scientific and Technological Development, Brazil). For the experiments conducted in Finland, this study was supported by European Research Council, grant #322742 - iPLASTICITY; EU Joint Programme - Neurodegenerative Disease Research (JPND) CircProt, project #301225 and #643417, Sigrid Jusélius Foundation, Jane and Aatos Erkko Foundation, and the Academy of Finland grants #294710 and #307416.

## Acknowledgment

The authors thank Flavia Salata (University of São Paulo) and Sulo Kolehmainen (University of Helsinki) for their technical assistance.

## Authors’ contribution

ISM designed study, performed the experiments, collected, analyzed and interpreted data, and wrote the manuscript draft. AJS performed the experiments, collected, analyzed data. PCC, EC, CB and SRLJ designed study, analyzed and interpreted data, and wrote the manuscript.

## Notes

### Competing Interest Statement

The authors have declared no competing interest.

